# Unique trajectory of gene family evolution from genomic analysis of nearly all known species in an ancient yeast lineage

**DOI:** 10.1101/2024.06.05.597512

**Authors:** Bo Feng, Yonglin Li, Hongyue Liu, Jacob L. Steenwyk, Kyle T. David, Xiaolin Tian, Biyang Xu, Carla Gonçalves, Dana A. Opulente, Abigail L. LaBella, Marie-Claire Harrison, John F. Wolters, Shengyuan Shao, Zhaohao Chen, Kaitlin J. Fisher, Marizeth Groenewald, Chris Todd Hittinger, Xing-Xing Shen, Antonis Rokas, Xiaofan Zhou, Yuanning Li

**Affiliations:** Institute of Marine Science and Technology, Shandong University, Qingdao 266237, China; Laboratory for Marine Biology and Biotechnology, Qingdao Marine Science and Technology Center, Qingdao 266237, China; Guangdong Laboratory for Lingnan Modern Agriculture, Guangdong Province Key Laboratory of Microbial Signals and Disease Control, Integrative Microbiology Research Center, South China Agricultural University, Guangzhou 510642, China; Howards Hughes Medical Institute and the Department of Molecular and Cell Biology, University of California, Berkeley, Berkeley, CA, USA; Department of Biological Sciences, Vanderbilt University, Nashville, TN 37235, USA; Evolutionary Studies Initiative, Vanderbilt University, Nashville, TN 37235, USA; Associate Laboratory i4HB—Institute for Health and Bioeconomy and UCIBIO— Applied Molecular Biosciences Unit, Department of Life Sciences, NOVA School of Science and Technology, Universidade NOVA de Lisboa, Caparica, Portugal; UCIBIO-i4HB, Departamento de Ciências da Vida, Faculdade de Ciências e Tecnologia, Universidade Nova de Lisboa, Caparica, Portugal; Laboratory of Genetics, J. F. Crow Institute for the Study of Evolution, Center for Genomic Science Innovation, Department of Energy (DOE) Great Lakes Bioenergy Research Center, Wisconsin Energy Institute, University of Wisconsin-Madison, Madison, WI 53726, USA; Biology Department, Villanova University, Villanova, PA 19085, USA; Department of Bioinformatics and Genomics, University of North Carolina at Charlotte, North Carolina Research Campus, Kannapolis NC 28223, USA AND Center for Computational Intelligence to Predict Health and Environmental Risks (CIPHER), University of North Carolina at Charlotte, Charlotte, NC, 28233, USA; Department of Biological Sciences, State University of New York at Oswego, Oswego, NY, 13126, USA; Westerdijk Fungal Biodiversity Institute, 3584 Utrecht, The Netherlands; Key Laboratory of Biology of Crop Pathogens and Insects of Zhejiang Province, Institute of Insect Sciences, Zhejiang University, Hangzhou 310058, China

## Abstract

Gene gains and losses are a major driver of genome evolution; their precise characterization can provide insights into the origin and diversification of major lineages. Here, we examined gene family evolution of 1,154 genomes from nearly all known species in the medically and technologically important yeast subphylum Saccharomycotina. We found that yeast gene family and genome evolution are distinct from plants, animals, and filamentous ascomycetes and are characterized by small genome sizes and smaller gene numbers but larger gene family sizes. Faster-evolving lineages (FELs) in yeasts experienced significantly higher rates of gene losses—commensurate with a narrowing of metabolic niche breadth—but higher speciation rates than their slower-evolving sister lineages (SELs). Gene families most often lost are those involved in mRNA splicing, carbohydrate metabolism, and cell division and are likely associated with intron loss, metabolic breadth, and non-canonical cell cycle processes. Our results highlight the significant role of gene family contractions in the evolution of yeast metabolism, genome function, and speciation, and suggest that gene family evolutionary trajectories have differed markedly across major eukaryotic lineages.

## Introduction

Gene duplications and losses are one of the major drivers of genome evolution and the source of major evolutionary innovations. For example, the evolutionary transition to vascular plants, originating from the common ancestor of Viridiplantae, was characterized by significant gene family expansion events, reflecting adaptations to life in terrestrial environments^1^. Similarly, the evolution of animals was marked by the accumulation of genes essential for multicellularity^2^. In contrast, the ancestors of fungi primarily experienced a reduction in most functional gene categories, with early fungal evolution featuring both the loss of ancient protist gene families and the expansion of novel fungal gene families^3^. These distinct evolutionary trajectories underscore the diversity and adaptive strategies of eukaryotes.

The Saccharomycotina subphylum (phylum Ascomycota, Kingdom Fungi) encompasses a diverse array of ∼1,200 species, including the well-known baker’s yeast *Saccharomyces cerevisiae*, the opportunistic pathogen *Candida albicans*, and the industrial producer of oleochemicals *Yarrowia lipolytica*^4,5^. Species in the subphylum, which began diversifying approximately 400 million years ago, showcase remarkable ecological, genomic, and metabolic diversity^6–11^. From fermenting sugars to metabolizing urea and xenobiotic compounds, yeasts have evolved diverse metabolic pathways that allow them to thrive in environments as varied as fruit skins, deep-sea vents, arctic ice, and desert sands^11–16^. Genome-wide protein sequence divergence levels within the yeast subphylum are on par with those observed within the plant and animal kingdoms^17^. However, gene family evolution in the yeast subphylum remains largely unexplored. This limitation has been primarily due to a concentration of research on a limited subset of species and the lack of comprehensive genomic data across the whole subphylum^18–20^. Moreover, evolutionary analyses of a wide range of yeast species would facilitate better understanding of the specific genes and genetic mechanisms enabling them to thrive in various ecological niches.

Here, we leveraged the recent availability of 1,154 draft genomes from 1,051 yeast species, covering 95% of known species within the Saccharomycotina subphylum, to investigate the relationship between gene family evolution and yeast diversity. Comparative analysis with three other major eukaryotic lineages—plants, animals, and filamentous ascomycetes—reveals that yeasts have smaller weighted average gene family sizes due to fewer gene counts. However, at similar gene counts, such as when comparing the yeast *Dipodascus armillariae* with 9,561 genes and the green alga *Micromonas pusilla* with 10,238 genes, yeasts exhibit larger weighted average gene family sizes (1.68 vs. 1.35 genes / gene family, respectively). Within three specific yeast taxonomic orders, we identified marked weighted average gene family size differences among distinct lineages that enabled us to categorize them into two distinct groups: faster-evolving lineages (FELs) characterized by faster rates of protein sequence evolution, higher numbers of gene family reductions and losses, and higher speciation rates; and slower-evolving lineages (SELs) that exhibited the converse pattern. The affected gene families are predominantly involved in key processes such as mRNA splicing, cell division, and metabolism. These changes, including the loss of introns and reduced diversity in carbon source utilization, suggest that dynamic gene family alterations, especially contractions, may have been key in shaping the evolutionary trajectory of yeast genomic and phenotypic diversity. Our findings underscore the significant impact of gene family dynamics on yeast evolution, revealing that contractions in gene families have resulted in fewer gene counts than filamentous ascomycetes, animals, and plants. Yet, yeasts have maintained higher weighted average gene family sizes than animals and filamentous ascomycetes. This finding provides both broad and fine-scale resolution of the tempo and mode of yeast evolutionary diversification.

## Results

### Gene Family Diversity is Correlated with Total Gene Content in Eukaryotes

We sampled 1,154 yeast genomes, 761 filamentous ascomycetous (from subphylum Pezizomycotina) genomes, 83 animal (Kingdom Metazoa) genomes, and 1,178 plant (Kingdom Viridiplantae, Phylum Glaucophyta, and Phylum Rhodophyta) genomes and transcriptomes from previous studies^1,11,21,22^, representing every major lineage across these four groups (Table S1). Using OrthoFinder, we identified 62,643 orthologous groups of genes (hereafter referred to as gene families) in yeasts, 137,783 in Pezizomycotina, 65,811 in animals, and 52,956 in plants. To filter out species-specific or rare gene families, we excluded all gene families that were present in 10% or fewer of the taxa in each major lineage (the threshold of 10% was based on the density plot of gene family average coverage; Figure S1). This filtering resulted in the identification of 5,551 gene families in yeasts (that collectively contain 89.88% of the genes assigned to orthogroups by OrthoFinder), 9,473 in Pezizomycotina (∼87.09%), 11,076 in animals (∼76.68%), and 8,231 in plants (∼96.41%).

Examination of weighted average gene family sizes, calculated using the reciprocal of maximum observed gene family size as the weight to account for differences in gene family size, revealed distinct features of gene family content for each group. Specifically, yeasts and filamentous ascomycetes typically had smaller weighted average gene family sizes than animals and plants (Figure 1a). However, when comparing organisms with equivalent numbers of protein-coding genes, yeasts displayed similar weighted average sizes to plants and larger sizes than filamentous ascomycetes and animals (Figure 1b).

**Figure 1:**
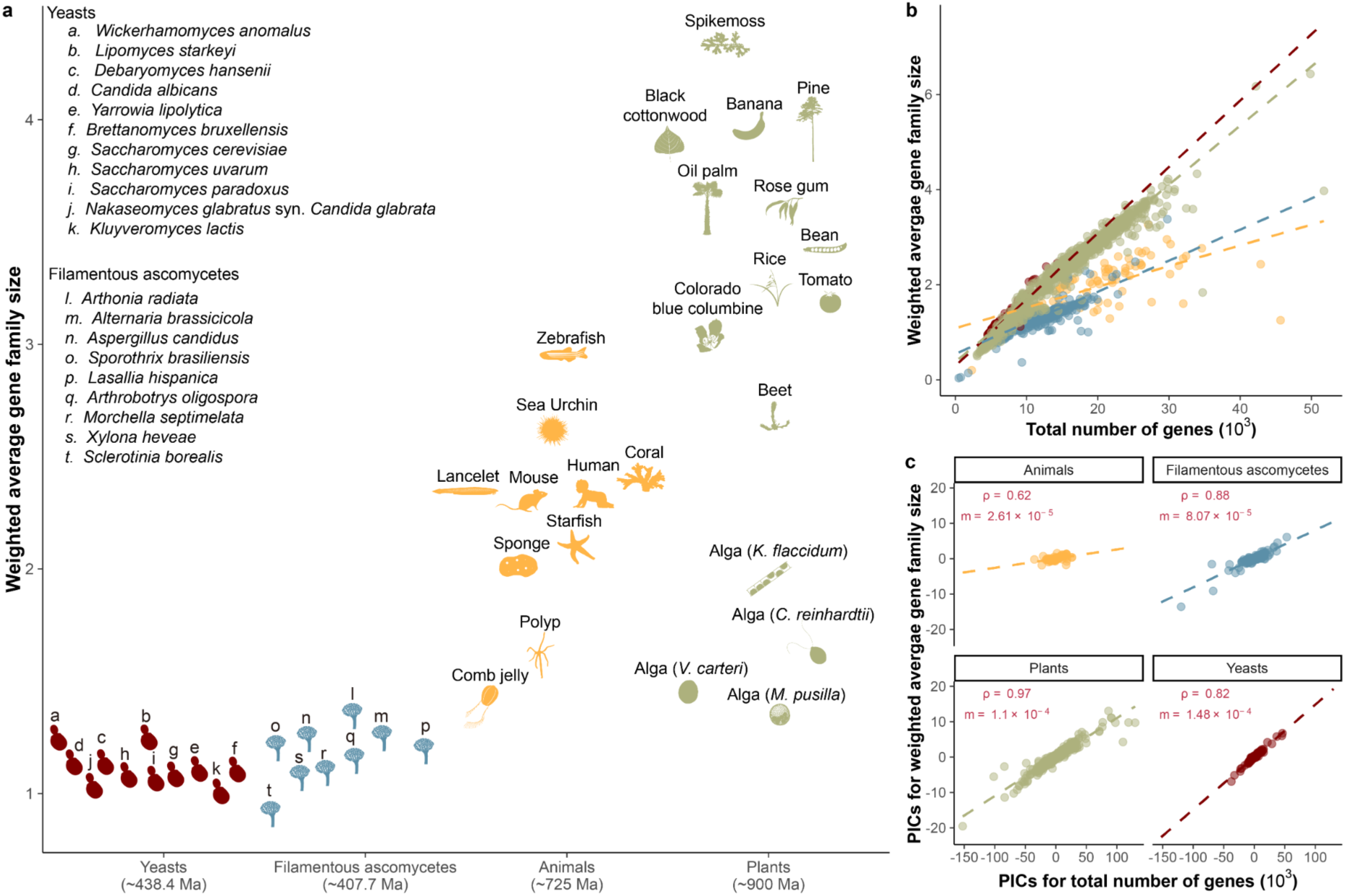
Narrow range of weighted average gene family sizes among yeasts versus broader diversity in animals and plants. a. The weighted average size of gene families across yeasts (from subphylum Saccharomycotina), filamentous ascomycetes (subphylum Pezizomycotina), animals (Kingdom Metazoa), and plants (Kingdom Viridiplantae, Phylum Glaucophyta, and Phylum Rhodophyta). Species-specific gene families were excluded by applying a 0.1 threshold based on the density plot for gene family average coverages (Figure S1). Representative species for yeasts and animals were identified based on previous studies^17^; representatives for plants were chosen from species with available genome data; for filamentous ascomycetes, one representative per class was selected. The estimated divergence times are approximately 438.4 million years for yeasts, 407.7 million years for filamentous ascomycetes, 725 million years for animals, and 900 million years for plants, derived from previous studies^17,21,73,74^. Images representing taxa were manually created and sourced from Phylopic (https://www.phylopic.org/). b. Correlation plot between the weighted average gene family size and the total number of protein-coding genes across yeasts, filamentous ascomycetes, animals, and plants. c. Correlation plot between the PICs of weighted average gene family size and the total number of protein-coding genes across yeasts, filamentous ascomycetes, animals, and plants. Correlations were determined through the Spearman test using the R package stats version 4.3.2. Specifically, the correlation coefficient (rho) for yeasts was 0.82, for filamentous ascomycetes was 0.88, for animals was 0.62, and for plants was 0.97, all statistically significant with *P* < 0.01. The slope (m) is calculated using linear regression based on the PICs of weighted average gene family size and the total number of protein-coding genes across these four groups. The PIC-related codes and data are available at the Figshare repository.

Moreover, we found a strong positive correlation between the phylogenetic independent contrasts (PICs) of weighted average gene family size and the number of protein-coding genes (gene number). This correlation was particularly pronounced in plants (rho = 0.97), yeasts (rho = 0.82), and filamentous ascomycetes (rho = 0.88), but weaker in animals (rho = 0.62), with all P-values less than 0.01 (Figure 1c and Table S2). The correlation between PICs of weighted average gene family size and genome size was weaker (Table S2). Our PIC regression showed yeasts had a steeper slope than plants, animals or filamentous ascomycetes (Figure 1c). This indicates that yeasts tend to have larger gene family sizes as their gene number increases (Figure 1b). This result suggests that yeasts tend to exhibit larger gene family sizes / gene number compared to animals and filamentous ascomycetes and are on par with plants, corroborating the contributions of gene duplications to yeast phenotypic diversity^23–25^.

### Reduced Gene Family Content is Associated with Rapid Genome Sequence Evolution

The weighted average gene family size across 12 yeast orders^26^ is 1.12 genes / gene family, with Alloascoideales having the highest size at 1.49 and Saccharomycodales having the lowest size at 0.82 (Figure 2a). The average gene number and genome size across all 12 orders is 5,908 genes and 13.17 Mb, respectively. Alloascoideales yeasts have the highest average gene numbers and genome sizes (8,732 genes and 24.15 Mb, respectively), whereas Saccharomycodales have the smallest ones (4,566 genes and 9.82 Mb, respectively).

**Figure 2:**
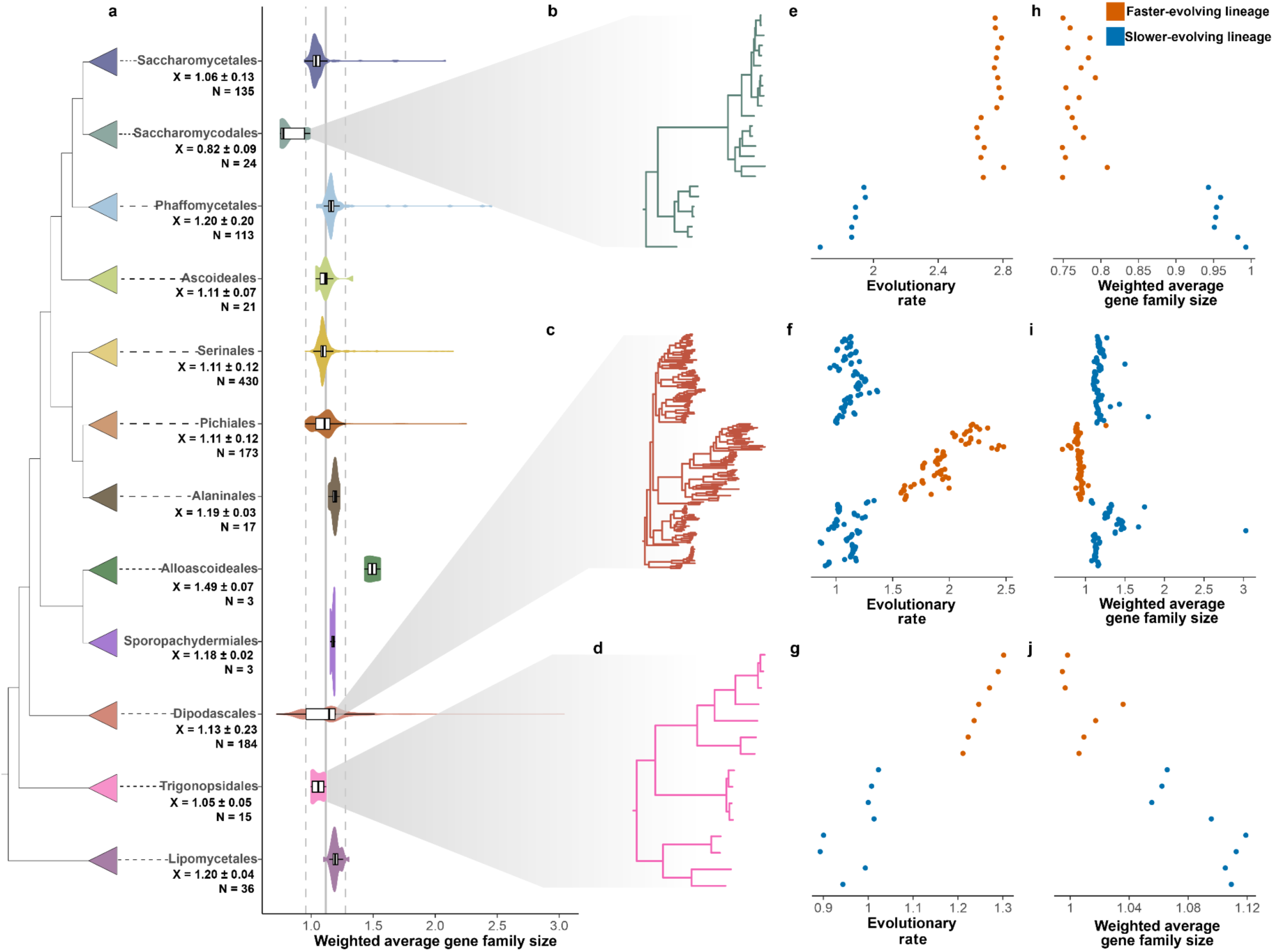
Notable variations in weighted average gene family sizes within specific yeast orders. a. The phylogeny of 1,154 yeasts, derived from a previous study^11^. Colors indicate the taxonomic classification of species within the Saccharomycotina order. The weighted average gene family sizes (X) and genome numbers (N) for each order are displayed beneath the respective order names. A gray solid line at 1.12 represents the weighted average gene family size for all yeasts. b-d. The orders Trigonopsidales, Dipodascales, and Saccharomycodales are highlighted due to their notable differences in evolutionary rates and weighted average gene family sizes. e-j. Differences in evolutionary rates / weighted average gene family sizes within specific orders. Each dot represents a yeast in the corresponding phylogeny and is arranged according to its placement on the phylogenetic tree.

Saccharomycodales contains the FEL in the genus *Hanseniaspora*, which is known to have experienced significant lineage-specific gene losses, especially in genes involved in the cell cycle and DNA repair, which are correlated with significantly higher evolutionary rates^27^. Thus, we first examined the correlation between weighted average gene family size and evolutionary rate across the 12 orders and found that it was moderate (rho = –0.41, *P* < 0.01) (Figure S3a). We next tested whether weighted average gene family size and evolutionary rate varied within specific orders. We found lineage-specific variations in evolutionary rates for Dipodascales (*P* = 0.04), Saccharomycodales (*P* = 0.01), Trigonopsidales (*P* < 0.01), Pichiales (*P* < 0.01), and Serinales (*P* < 0.01) using the multimodality test (Table S3). However, only Dipodascales, Saccharomycodales, and Trigonopsidales showed lineage-specific variations in their weighted average gene family sizes (Figures 2b-j and S4). Examining the relationship between weighted average gene family size and evolutionary rate uncovered two distinct clusters within each order (Figures 2b-j and S5). These clusters corresponded to faster-evolving lineages (FELs), characterized by smaller weighted average gene family sizes and higher evolutionary rates, and slower-evolving lineages (SELs), which exhibited larger weighted average gene family sizes and slower evolutionary rates. Specifically, differences in weighted average gene family size included median values of genes / gene family of 1.01 for FEL vs. 1.10 for SEL in Trigonopsidales, 0.93 vs. 1.17 in Dipodascales, and 0.76 vs. 0.95 in Saccharomycodales (all *P* < 0.01). For evolutionary rates, the average number of amino acid substitutions / site were 1.25 vs. 1.00 in Trigonopsidales FEL vs. SEL, 1.93 vs. 1.12 in Dipodascales FEL vs. SEL, and 2.75 vs. 1.89 in Saccharomycodales FEL vs. SEL (all *P* < 0.01). Notably, all three FELs formed clades that were distinct from or emerged within SELs on the yeast phylogeny (Figures 2b-d) and significantly differed in their speciation rates from SELs in two of the three lineages (DR statistic median of 0.03 vs. 0.02 in Dipodascales FEL vs. SEL, *P* < 0.01; 0.12 vs. 0.02 in Saccharomycodales FEL vs. SEL, *P* < 0.01; 0.01 vs. 0.01 in Trigonopsidales FEL vs. SEL, *P* = 0.27) (Figure 3e).

**Figure 3:**
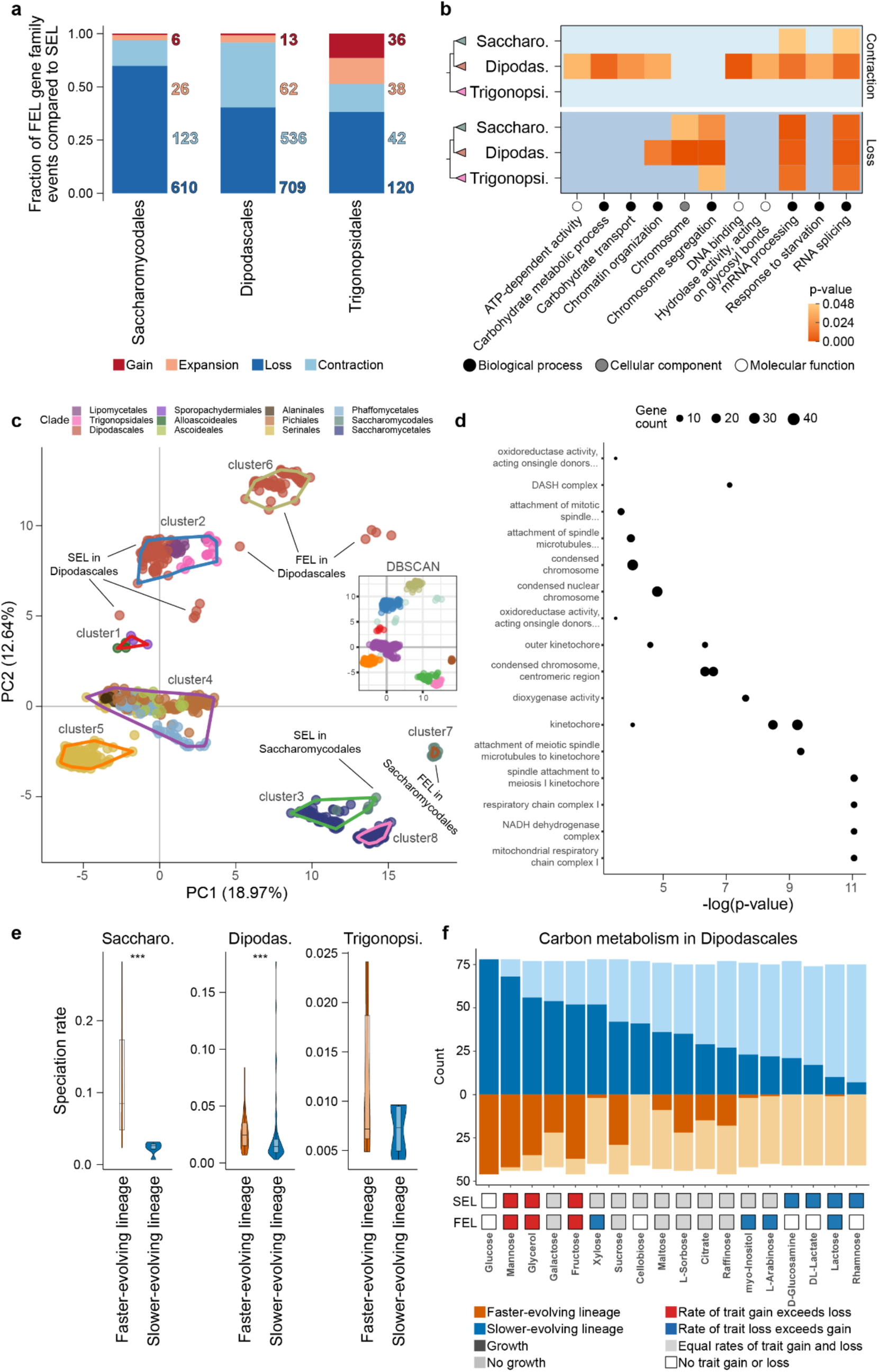
Faster-evolving lineages (FELs) within three orders experienced significantly more gene family contractions and losses. a. Significantly different gene family dynamics (loss, contraction, expansion, and gain) in FELs relative to SELs within Dipodascales, Saccharomycodales, and Trigonopsidales. A gene family loss is indicated by a fold change value of 0, meaning the gene family in FEL has no copies, while a fold change equal to positive infinity signifies gain. Values greater than 1.5 indicate expansion, and values less than 0.67 signify contraction. The Kolmogorov–Smirnov test was employed to assess these differences; *P* ≤ 0.05. b. GO enrichment analysis of significant contractions or losses in gene families. All enriched GO terms were simplified into GO slim terms. c. PCA analysis utilizing presence and absence data for 4,262 gene families with an average coverage of 0.5 or greater. The DBSCAN plot employs PC1 and PC2 coordinates for density-based clustering, with colors distinguishing the various clusters. In the PCA plot, points enclosed by lines indicate distinct clusters, corresponding to the color coding applied in the DBSCAN plot. d. The GO enrichment analysis of the top 610 gene families from PC1. e. Speciation rate comparison between FEL and SEL within Trigonopsidales, Dipodascales, and Saccharomycodales with the Wilcoxon signed-rank test. f. The evolutionary history of 17 carbon traits in FEL and SEL of Dipodascales. The dark color indicates the number of yeasts capable of utilizing the carbon source. Three different evolutionary models are shown: trait gain (red), trait loss (blue), and equal rates of trait gain and loss (gray). Estimated evolutionary models were not derived for glucose in both FEL and SEL, and for cellobiose, D-glucosamine, DL-lactate, and rhamnose in SEL, due to the uniform ability or inability of all yeasts within the group to utilize these carbon sources.

To identify gene families with significantly different sizes between FELs and SELs, we examined the fold change in average size (non-weighted) for each gene family and for each pair. Following a previous study^1^, we categorized changes into loss events (fold change equal to 0 in FEL vs. SEL), contractions (fold change < 0.67 in FEL vs. SEL), expansions (fold change > 1.5 in FEL vs. SEL), and gains (fold change ∼infinity in FEL vs. SEL). We found extensive and significant gene family losses and contractions in FELs (adjusted *P* ≤ 0.05) (Figures 2k-m). Specifically, the fractions of gene families that experienced significant contraction or loss in FELs were 10.40% (536/5,155) and 13.75% (709/5,155) in Dipodascales, 3.03% (123/4,056) and 15.04% (610/4,056) in Saccharomycodales, and 0.89% (42/4,727) and 2.54% (120/4,727) in Trigonopsidales.

### Rapidly Evolving Lineages Lost Genes Related to RNA Splicing, Cell Division, and Metabolism

To determine the functions of gene families contracted or lost in FELs, we performed enrichment analyses using three annotation datasets—Gene Ontology (GO) terms, InterPro annotations, and Kyoto Encyclopedia of Genes and Genomes Ortholog (KO). Functional categories enriched among gene families significantly contracted or lost in FELs relative to SELs yielded numerous GO terms common across the three orders, including those associated with transcriptional functions, like RNA splicing and mRNA processing (Figure 3b). Additionally, the Dipodascales FEL experienced significant contractions in gene families related to carbohydrate metabolism. Our InterPro and Kyoto Encyclopedia of Genes and Genomes (KEGG) enrichment analyses confirmed these findings (Table S4).

In addition to comparing weighted average gene family size between FELs and SELs, we illustrated the differences among yeasts based on the presence (1) and absence (0) of gene families. To exclude outliers (species-specific and/or rare gene families), we set the threshold to 0.5 based on the bimodal distribution (Figure S1) and carried out all subsequent analyses. A more relaxed threshold of 0.1 gave rise to highly consistent PCA distribution and correlation results (Figures 3c and S6). Therefore, we discuss results from using the 0.5 threshold hereafter.

Following the PCA, density-based clustering according to the yeasts’ position on the first two principal components (PC1 and PC2) indicated that the distributions of clusters (each corresponding to one or a few orders) generally follow the phylogeny of these orders (Figures 2a and 3c), suggesting that patterns of gene presence or absence largely reflect yeast evolutionary relationships. Moreover, consistent with our previous findings from the fold change analysis, FELs and SELs were separated into two distinct clusters in Dipodascales (FEL in cluster 6, SEL in cluster 2) and Saccharomycodales (FEL in cluster 7, SEL in cluster 3). The FEL and SEL from Trigonopsidales were not segregated into distinct groups but were spaced apart in cluster 2. Notably, all 3 of these orders showed significant differences in the PC1 coordinates between FELs and SELs (*P* ≤ 0.05) (Figure S7).

To determine which gene families’ presences or absences contribute to the distribution variation among yeasts in the PCA scatter plot, we investigated the correlation between the presence or absence of yeast gene families and their coordinates on the principal components. We identified 610 gene families whose average presence and absence in yeasts were most strongly correlated with their PC1 coordinates (rho = –0.99, *P* < 0.01), explaining significant species variation along this axis (Figure 4c). The strong negative correlation indicates that an increase in PC1 coordinates correlates with losses in the 610 gene families, with Saccharomycodales, Saccharomycetales, and the FEL from Dipodascales experiencing more losses than other lineages (Figures 3c and S8). In contrast, there was no clear relationship for gene family presence or absence along PC2 (Figure S9). We employed the same enrichment analysis method used in the fold change analysis on these 610 gene families, revealing GO terms related to oxidoreductase activity; mitochondrial electron transport chain; and notably, cell division processes, such as the kinetochore, condensed chromosome, and DASH complex (Figure 3d). Our InterPro and KEGG analyses echoed these findings (Table S5). The enrichment results from both the fold change analysis and PCA analysis of gene presence/absence pattern (PCA analysis for short afterwards) highlighted GO terms associated with meiotic processes (adjusted *P* ≤ 0.05). These include meiotic chromosome segregation (GO:0045132), kinetochore (GO:0000776), and the attachment of meiotic spindle microtubules to kinetochore (GO:0051316).

**Figure 4:**
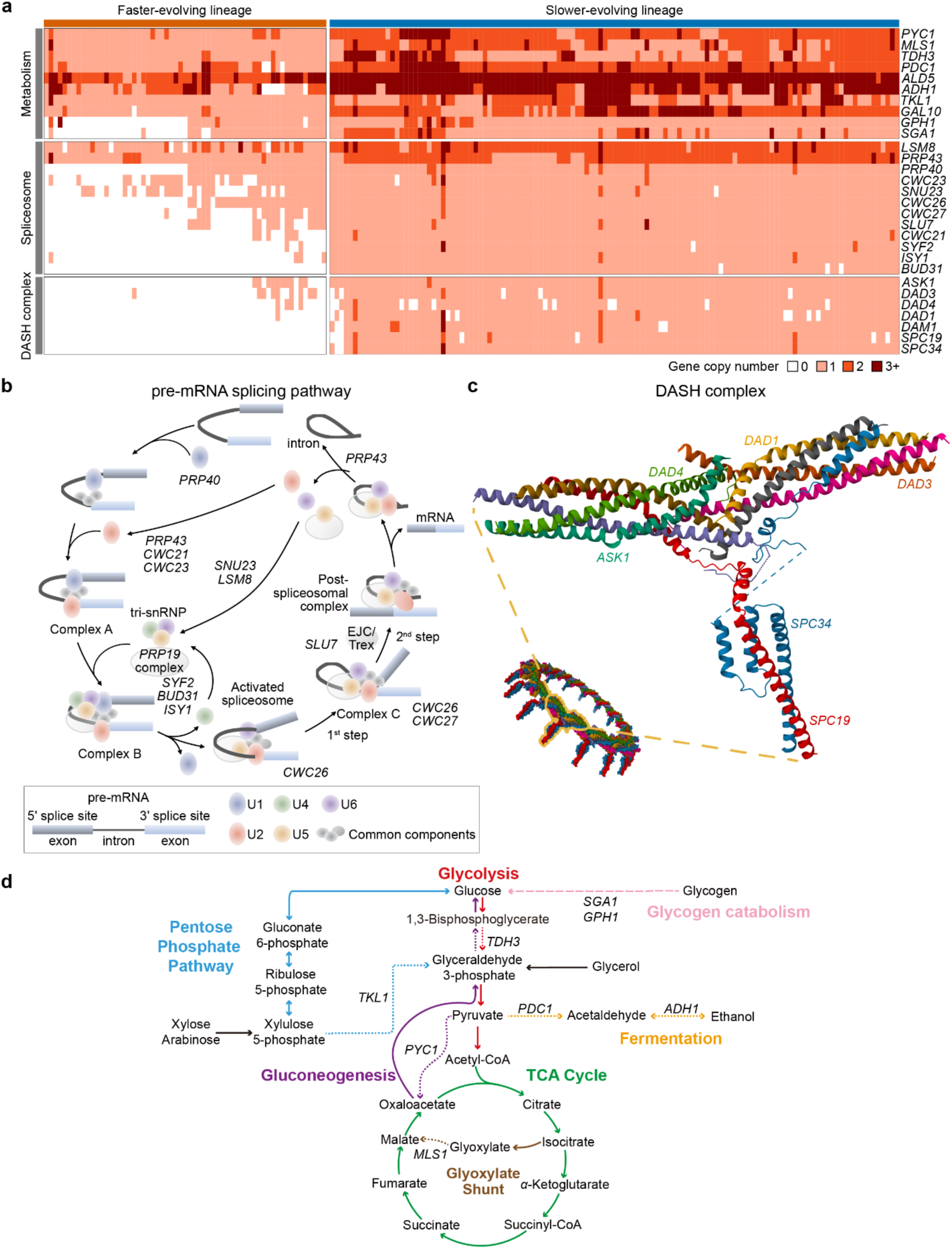
Dipodascales’ FEL experienced the loss of key genes involved in the pre-mRNA splicing pathway, metabolic pathways, and the DASH complex. a. A detailed picture of gene copy numbers in Dipodascales among metabolic pathways (10 gene families), the pre-mRNA splicing pathway (12 gene families), and the DASH complex (7 gene families). Column colors indicate SEL (yellow) and FEL (green). The estimated gene family names, identified using *S. cerevisiae* as a reference, are listed to the right of the columns. b. The pre-mRNA splicing pathway. Gene family names are marked at specific steps encoded in the pathway that experienced contractions or losses in the FEL. c. Genes encoding the DASH complex. d. Carbon metabolism pathways containing widespread gene loss or contraction in the Dipodascales FEL. Pathway names and reactions are indicated in corresponding colors. Steps encoded by genes experiencing contraction or loss are represented by dashed lines labeled with the gene name (gene family contractions – short dashes, gene family losses – long dashes). Pathways are abridged to show steps relevant to reported losses and contractions and not all intermediate metabolites are shown. Black arrows indicate where glycerol (gained in FEL) and xylose & arabinose (lost in FEL) feed into central carbon metabolism.

### Gene Family Losses Suggest Non-canonical Spliceosomes, Metabolic Pathways, and DASH Complexes within the FEL of Dipodascales

To explore which gene families and pathways—within the enriched functional categories—experienced contraction or loss in FELs, we mapped gene families enriched in the fold change analysis and PCA analysis to the KEGG database and *Saccharomyces* Genome Database (SGD)^28^, using the *S. cerevisiae* genome as a reference. Given that the FEL in Dipodascales exhibited the most significant contractions and losses of gene families compared to Saccharomycodales and Trigonopsidales, and the enrichment of RNA splicing, the DASH complex and metabolic process in fold change or PCA analyses, our study concentrated on Dipodascales. In terms of functions, we focused on the pre-mRNA splicing pathway, metabolic pathways, and the DASH complex.

The pre-mRNA splicing pathway primarily removes introns from pre-mRNA and joins exons, forming mature mRNA for protein synthesis^29^. In this pathway, 14% of the genes (12/85) exhibited contractions or losses. While *LSM8* and *PRP43* significantly contracted in the Dipodascales FEL, other gene families experienced extensive losses (Figure 4a and b). These include *PRP40*, *CWC21*, *SNU23*, and *CWC23*, which are associated with the assembly of the spliceosomal subunits U1, U2, U4, U5, and U6^29^. Almost all species in the Dipodascales FEL have lost genes related to the Prp19 complex, which is crucial for promoting the assembly and activation of the spliceosome, as well as stabilizing its structure^30^. These losses could ultimately lead to abnormalities in splicing mechanisms. Notably, we found that there was significant intron loss in the Dipodascales FEL both in the total number of introns (TNI) and the average number of introns per gene (ANI) within species, with a stark reduction from a median TNI of 2,815 per SEL species to 466 per FEL species (*P* < 0.01) and a decrease in ANI from 1.44 to 1.31 (*P* < 0.01) (Figures S10b and c). Similar pattern of significant intron loss was observed in Trigonopsidales, with a median TNI of 6,287 per SEL species vs. 789 per FEL species (*P* < 0.01) and a median ANI of 2.05 per SEL species vs. 1.31 per FEL species (*P* < 0.01) (Figures S10b and c). In Saccharomycodales, the pattern was more subtle, with a median TNI of 528 per SEL species vs. 252 per FEL species (*P* = 0.01) and a median ANI of 1.22 per SEL species vs. 1.20 per FEL species (*P* = 0.29) (Figures S10b and c).

The DASH complex plays a crucial role in eukaryotic cell division, particularly in chromosome segregation during mitosis^31^. Strikingly, genes associated with the DASH complex were extensively lost in the Dipodascales FEL, such as *ASK1*, *DAD3*, *DAD4*, and *DAD1*, which are integral components of this complex (Figure 4c). *DAM1*, *SPC19*, and *SPC34* were lost entirely in Dipodascales FEL species. The loss of *DAM1*, primarily involved in the stability of kinetochore microtubules, likely results in compromised microtubule stability^32^. Similarly, the absence of *SPC19* and *SPC34*, critical for the attachment of the kinetochore to microtubules, potentially leading to defects in chromosome segregation^33^.

Key metabolic pathways also exhibited considerable variation in gene family size in the Dipodascales FEL. More than half of these yeasts have lost *GPH1* and *SGA1* in the carbohydrate degradation pathway, which are responsible for encoding glycogen phosphorylase and sporulation-specific glucoamylase, respectively (Figures 4a and d). The loss of *GPH1* and *SGA1* genes likely affects Dipodascales FEL’s ability to utilize glycogen and amylopectin-like polysaccharides^34,35^. Furthermore, significant contractions were observed for *MLS1*, which encodes a key step in the glyoxylate shunt of the TCA cycle; *PYC1,* which encodes the enzyme that converts pyruvate to oxaloacetate where it can enter the TCA cycle or gluconeogenesis; *PDC1*, *ADH1*, and *ALD5*, which encode key steps in fermentation; and *TKL1*, which encodes two key reactions in the pentose phosphate pathway. We note that the present analyses reflect the known loss of the *PDC1* and *ADH1* genes in several members of the *Wickerhamiella*/*Starmerella* (W/S) clade of the Dipodascales FEL^36^, but many of them reacquired alcoholic fermentation through the horizontal transfer of bacterial genes encoding alcohol dehydrogenases and the cooption of paralogs encoding decarboxylases. Further, a single FEL clade of 4 *Starmerella* species has lost *PCK1* and *FBP1*, genes essential for gluconeogenesis, *ICL1*, which encodes an essential component of the glyoxylate shunt, *GSY1*, which encodes glycogen synthase, and *GPH1* and *GDB1,* which encode the glycogen phosphorylase and glycogen debranching enzymes required for degradation of glycogen. Complete loss of *PCK1* and *FBP1* in a free-living yeast has previously been reported only in the Saccharomycodales^27^.

For gene families that experienced significant contractions or losses in the pre-mRNA splicing pathway, metabolic pathways, and the DASH complex in Dipodascales FEL, we observed consistent, but less pronounced, patterns in Saccharomycodales and Trigonopsidales FELs. Specifically, in the pre-mRNA splicing pathway, 50% (6/12) of genes displayed significant losses in fold change analysis in Saccharomycodales, while Trigonopsidales showed no significant changes in these genes (Table S6). All genes in Saccharomycodales had significant losses for the DASH complex, with only *DAD1* and *SPC19* similarly affected in Trigonopsidales (Table S6). No significant results were found in the metabolic pathways for genes lost in Dipodascales for either Saccharomycodales or Trigonopsidales. This outcome aligns with our enrichment results, where only a few GO terms related to these functions were enriched in Trigonopsidales, and metabolic-related functions were predominantly enriched in Dipodascales (Figure 3b and d).

To investigate potential impacts on carbon source utilization in Dipodascales FEL, we analyzed the evolutionary trends of 18 major carbon sources^11^. We found a distinct tendency for FEL to lose growth traits associated with these carbon sources (Figure 3f). For instance, while SEL species retained the ability to utilize cellobiose, D-glucosamine, DL-lactate, and rhamnose, FEL species have lost these growth traits. Furthermore, we found that the rate of acquiring xylose, myo-inositol, and L-arabinose growth traits in SEL species was equal to the rate of losing them.

However, in FEL species, the loss rate surpassed the gain rate. Interestingly, both FEL and SEL species exhibited a greater tendency to acquire the glycerol growth trait, despite the *TDH3* gene family, which is crucial for glycerol metabolism (as well as glycolysis and gluconeogenesis), has undergone significant contraction in FEL. This result suggests the possibility of other genes or pathways being augmented to compensate for the *TDH3* contraction and enable glycerol metabolism^37^. These observations suggest that gene losses and contractions in Dipodascales FEL species have significantly altered their metabolic capacities.

### Some Functional Categories Undergo Waves of Gains and Losses

Ancestral reconstructions of gene family content revealed waves of gains and losses, with a general trend of net gene loss from the Saccharomycotina common ancestor (SCA) to the most recent common ancestor (MRCA) of each order (tips in the Figure 5, hereafter only use order names instead). The exception was Dipodascales, which experienced a net gain of 543 genes. Certain nodes underwent notable changes in gene number; for instance, ancestral nodes such as <15>, Lipomycetales, and Trigonopsidales lost over 1,000 genes each, whereas the Alloascoideales, Dipodascales, Phaffomycetales, Pichiales, Serinales, and Saccharomycetales ancestors gained over 1,000 genes each.

**Figure 5:**
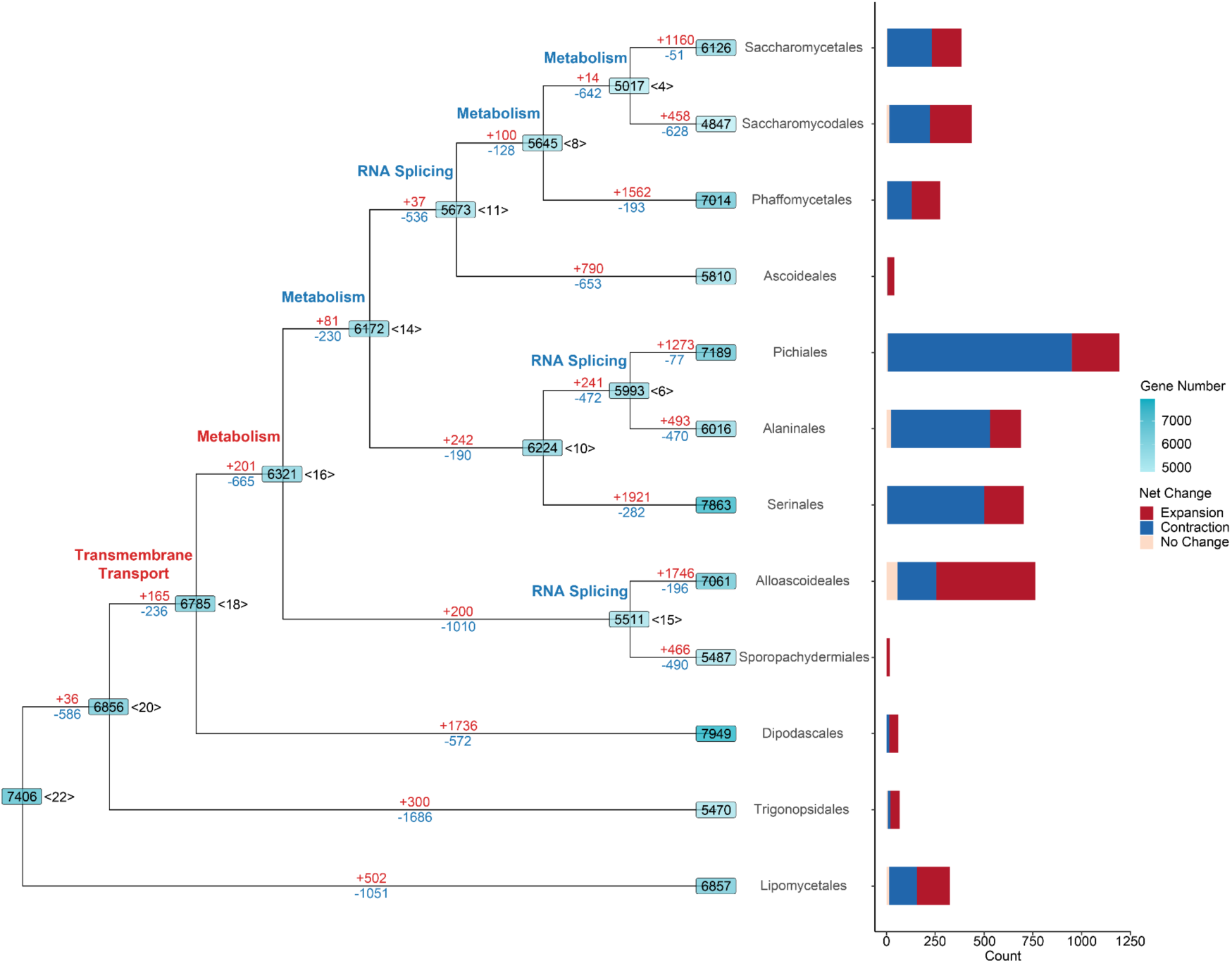
Yeasts have undergone a complex evolutionary history of gene families. The branches following the MRCA of each order have been collapsed to simplify the tree structure. Gene counts are marked on each node, with the corresponding node label positioned to its right. Gene gains are highlighted in red, while losses are depicted in blue along each branch. Additionally, branches are annotated with key terms from enriched GO terms (*P* ≤ 0.05); here, red signifies gene family expansion, and blue denotes contraction. A bar plot to the right of the tree quantifies the net changes in gene families within the phylogeny after the MRCA of each order. The y-axis, labeled “count”, reflects the number of gene families that underwent net changes—categorized into expansion, contraction, or no change. Expansion of a gene family is defined by a sum of net changes in copy number across all branches of an order being greater than 0, while contraction is defined by a sum less than 0, and no change is defined as a net change equal to 0.

Gene families within functional categories highlighted in previous analyses showed significant contractions and losses at ancestral yeast nodes. Specifically, gene families related to RNA splicing underwent substantial contractions at ancestral nodes <6>, <11>, <15>, Lipomycetales and Trigonopsidales, while expansions were observed at ancestral nodes Alaninales and Trigonopsidales (Figure 5 and Table S7a). Gene families involved in metabolism experienced frequent shifts, with contractions at ancestral nodes <4>, <8>, <14>, <16>, Ascoideales, Lipomycetales, Saccharomycetales, Saccharomycodales, Sporopachydermiales and Trigonopsidales (Figure 5 and Table S7a). Conversely, expansions were observed at ancestral nodes <4>, <16>, <18>, <20>, Alaninales, Alloascoideales, Lipomycetales, Serinales, and Trigonopsidales (Figure 5 and Table S7b). Gene families associated with transcription also exhibit a complex evolutionary history, showing contractions at ancestral nodes <14>, Ascoideales, Lipomycetales, Serinales, Sporopachydermiales, and Trigonopsidales, and expansions at ancestral nodes <4>, <16>, <18>, Alaninales, Alloascoideales, and Lipomycetales (Table S7b).

To investigate the evolutionary trends of gene families that experienced significant contractions or expansions in CAFE analyses within each yeast order, we calculated the net change of these gene families (net gain or loss across all branches). In orders that include Alaninales (508/689), Pichiales (943/1,194) and Serinales (498/704), over 70% of gene families with net changes experienced contractions, while in Alloascoideales (507/762), 66% of the events were gene family expansions (Figure 5). The remaining orders exhibited a nearly balanced mix of gene family expansion and contraction events. Gene families with net expansions were enriched in plasma membrane and transmembrane transporter-related GO terms (Table S8b). Conversely, DNA polymerase activity was prevalent in some gene families undergoing contractions, except in Serinales and Trigonopsidales, which are enriched in ligase activity and DNA repair functions, respectively (Table S8a).

To explore novel genes gained in the most recent common ancestor of each order, we selected orphan gene families (i.e., order-specific gene families) as determined by the coverage of each gene family across each order. Examination of orphan genes revealed variation among orders. Alloascoideales, Specifically and Sporopachydermiales orders each possessed over 180 orphan gene families, while other orders had fewer than 80 (Figure S11). The Dipodascales and Trigonopsidales orders each had only two orphan gene families, while Pichiales had one. Orphan genes were not enriched in specific functional categories.

## Discussion

Examination of gene family evolution of 1,154 genomes of nearly all known Saccharomycotina species elucidated, for the first time ever, the landscape of gene family evolution across a eukaryotic subphylum. Reductive evolution emerges as the main theme, marked by a transformation from a versatile SCA to descendants with more specialized lifestyle/metabolic capacity^17^ and smaller gene repertoires (Figure 5). In extant species, most yeasts exhibited similar weighted average gene family sizes and evolutionary rates. However, significant differences were observed in FELs compared to their SEL relatives in several independent yeast orders. The gene family size differences between FELs and SELs, enriched in similar functional categories, suggest that the same evolutionary trajectory has occurred repeatedly and independently in multiple yeast orders, indicating a broader trend rather than isolated incidents. The FELs demonstrated notable contractions and losses in gene families, especially those related to RNA splicing and the DASH complex (Figure 3b and d). Alterations in the pre-mRNA splicing pathway could generate novel transcript variants, potentially allowing some yeasts to better respond to environmental changes^29,38^. Additionally, impairments in the DASH complex may cause genomic instability, which, although potentially harmful under stable conditions, might provide adaptive advantages in fluctuating environmental stresses by increasing genetic diversity^31,39^.

These gene family contractions and losses in FELs may contribute to their higher evolutionary and speciation rates (Figures 2e-f and 3e) by enabling rapid genomic adaptations that optimize cellular processes crucial for survival and reproduction in diverse and challenging environments. For example, the FEL of Dipodascales is primarily found in the Arthropoda environment^11^, which is partially characterized by the production of various antifungal compounds and generally hostile conditions for many microorganisms^40,41^. This lineage also shows significant contractions in gene families related to metabolism and a general loss of growth traits, with a notable exception being the acquisition of glycerol utilization abilities (Figure 3b and f). This capability could be a key adaptation allowing them to thrive in specialized environments. Interestingly, a similar adaptation has been observed in endosymbionts like *Buchnera aphidicola* in aphids and *Wigglesworthia glossinidia* in flies, both of which effectively utilize glycerol^42^. The expansion of cytochrome P450 and cytochrome c oxidase assembly protein subunit gene families in Saccharomycodales and Dipodascales FELs (Table S5) suggests enhanced detoxification and metabolism of xenobiotic compounds, supporting their adaptation to hostile environments^43–45^. CAFE analysis has shown that certain functional categories, such as RNA splicing, metabolism, and cytochrome P450, are affected at more ancestral nodes in the yeast phylogeny (Figure 5 and Tables S7a and b). This suggests that the similar evolutionary trajectory observed across multiple yeast orders may be influenced by reductive evolution throughout the evolutionary history of yeast.

Moreover, yeasts and filamentous ascomycetes typically have smaller weighted average gene family sizes than animals and plants (Figure 1a) due to the strong correlation between the PICs of the weighted average sizes and gene numbers among these four groups (Figure 1c). Several key whole genome duplication (WGD) events occurred at the base of the animal and plant phylogenies^1,46,47^. In contrast, only one such event is known to have occurred near the base of a yeast order, affecting a small portion of the yeast phylogeny^7^; however, numerous other instances of hybridization, which could potentially result in WGD, have been noted in Saccharomycotina yeasts^48^. Meanwhile, fungal genomes, particularly those of yeasts, have undergone streamlining throughout evolution^49–51^. Additionally, many ancestral branches of yeasts exhibited widespread net gene loss in the CAFE analysis (Figure 4). This streamlining may contribute to their lower gene counts and weighted average gene family sizes compared to plants and animals. However, when we control for gene number, we found that yeasts exhibit larger weighted average gene family sizes than both filamentous ascomycetes and animals and are on par with plants (Figure 1b). Larger gene family sizes may provide redundancy and adaptability in biological processes, potentially enhancing the metabolic and stress response capabilities of yeast species and allowing them to thrive across diverse environmental conditions^52^.

State-of-the-art evolutionary genomic and phylogenomic studies now routinely report or analyze genomic data from hundreds to thousands of genomes^1,11,21,53^, ushering us in the “Thousand Genomes Era”. Analyzing gene families across thousands of genomes presents substantial challenges, including handling large datasets, accurately identifying and comparing complex genomic variations, and offering detailed functional annotations for a diverse range of genes. Traditional gene family analyses often concentrate on specific gene families, species, and gene family size evolution, leading to a gap in large-scale comparative analysis.

In this study, we implemented a comprehensive approach to explore gene family size differences across and within yeast lineages. We leveraged calculations of weighted average gene family size, comparisons based on evolutionary rates, and statistical tests to uncover evolutionary relationships and significant changes in gene families. This approach categorized yeasts into groups for fold change statistical analyses, identifying gene family dynamics, such as expansions and contractions, and comparing these within different yeast lineages to understand evolutionary pressures and trajectories. Additionally, we analyzed gene family composition, correlating gene presence or absence with yeast species distribution and identifying key gene families contributing to observed patterns. To reconstruct the evolutionary history of gene families across over 1,000 genomes, we employed a two-step approach (first calculating ancestral states within each order and then between orders) for gene family size estimation in ancestral yeasts, enhancing this approach with a detailed pipeline for broad-scale analysis. Our findings highlight gene family dynamics, such as losses and contractions, and establish a comparative framework for analyzing gene families at a large scale that can be readily applied to other major branches of the tree of life.

## Methods

### Data Collection and Collation

For our study on gene family evolution within Saccharomycotina yeasts, we acquired a comprehensive dataset comprising 1,154 Saccharomycotina yeast genomes. In addition, 21 non-budding yeast species were sampled as outgroups based on current understanding of Ascomycota phylogeny. These genomes, along with their annotations and a species tree, were obtained from our previous study^11^. This dataset provides a robust foundation for examining the evolutionary dynamics of gene families in yeasts. To compare the tempo and mode of gene family evolution of yeasts to other major eukaryotic lineages, we expanded our dataset to include 761 filamentous ascomycetes (Pezizomycotina) genomes^21^, 1,178 plant genomes and transcriptomes^1^, and 83 animal genomes^22^, including gene annotations for each. For all genomes, we kept the amino acid sequence translated from the longest protein coding sequence (CDS) from each gene. For plant transcriptomes, we adopted a protocol from^1^, using cd-hit version 4.8.1^54^ with a 99% sequence identity threshold to minimize redundancy. NCBI taxonomy and source information of all genomes and transcriptomes included in this study are also provided in Table S1 and the Figshare repository. Saccharomycotina species names in the supplementary tables and the Figshare repository were the current species names at the time used in the recent study^11^. For synonymous names and recent taxonomic updates, we refer the reader to the online MycoBank database.

### Delineation of Gene Family and Functional Annotation

To infer a comprehensive profile of gene families in budding yeasts, we delineated groups of orthologous genes (orthogroups, hereafter referred to as gene families) for the Saccharomycotina yeast dataset using OrthoFinder version 3.0, with default settings^55^. Following the approach of previous studies^56–58^, we used orthogroups from OrthoFinder as gene families. For consistency, we applied the same method to categorize gene families in Pezizomycotina, animal, and plant datasets. Due to the large number of genomes and transcriptomes in plants, we initially processed protein sequences from 30 representative genomes with the “-core” parameter to establish base orthogroups, and subsequently classified the protein sequences from an additional 1148 transcriptomes using the “-assign” parameter.

To obtain functional information of yeast gene families, we annotated all yeast genes from three independent aspects, including InterPro protein domains, and Gene Ontology (GO) and Kyoto Encyclopedia of Genes and Genomes (KEGG) terms. InterPro annotations were generated using InterProScan as part of a previous study^11^. GO annotations were generated using the eggNOG-mapper version 2.1.9^59^ with the search mode set to “mmseqs”. We initially compared KEGG annotations using the web-based GhostKOALA version 2.0^60^ with the KofamKOALA based annotations used in the study^11^. Due to GhostKOALA providing annotations for a larger number of gene families, we ultimately chose to exclusively use GhostKOALA for our final KEGG annotations.

### Weighted Average Gene Family Size Analysis of Gene Family Evolution

To assess the variations in gene family size among yeasts, Pezizomycotina, animals, and plants, we calculated the weighted average gene family size using a custom R script according to the following formula described in^1^. The weighted average gene family size reflects the overall size of gene families in a set of species, taking into account the relative size of each gene family.

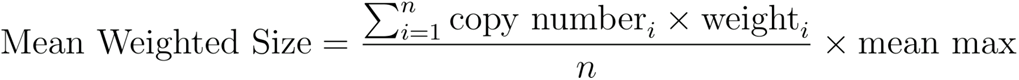

Taking yeasts as an example, in the formula, ‘n’ represents the total number of gene families in the dataset, ‘i’ stands for a specific gene family, and ‘weight’ is the reciprocal of the observed maximum copy number of the gene family among all yeast genomes. ‘Mean max’ is the average of the maximum copy numbers of these ‘n’ gene families.

Our preliminary analysis revealed a large number of gene families with highly restricted taxon distribution, which may confound the calculation of weighted average gene family size. Therefore, we implemented a lineage-based coverage assessment method^3^ for gene families across different taxa to exclude species-specific gene families. Specifically, we focused on assessing the coverage of each gene family within these 4 distinct groups, using yeasts as an example. Coverage in this context refers to the proportion of species within each clade that possesses a particular gene family. Using yeasts as an example, for each gene family, we first calculated its coverage in each of the 12 yeast orders^26^, and then took the average value as the overall coverage of the gene family. Similar procedures were followed for Pezizomycotina (9 classes), animals (14 phyla), and plants (22 phyla). Gene families with low average coverage values are likely to be highly species-specific. Given a bimodal distribution in the density plots of average coverage for gene families, we established a threshold of 0.1 to identify species-specific gene families (Figure S1). Families with average coverage below this threshold were considered species-specific for further analysis. This exclusion criterion was applied uniformly across the 4 groups studied.

To robustly test the correlation between weighted average gene family sizes and gene counts while accounting for phylogenetic relationships, we first converted the data into phylogenetic independent contrasts (PICs) using the “pic” function from the R package ape version 5.7.1, based on the respective phylogenetic trees for yeasts, filamentous ascomycetes, animals, and plants. Phylogenetic trees were obtained from previous studies^11,21,22^ and pruned to include only the species we studied using gotree version 0.4.4^61^. In the previous study^1^, the plant phylogenetic tree was constructed using ASTRAL, which is not optimized for accurate branch length estimation. Therefore, we retained the original tree topology and protein sequences from the previous study to reconstruct branch lengths using IQ-TREE version 2.2.3^62^. We then conducted a Spearman correlation test between these transformed datasets using the cor.test function (method = spearman) from the R package stats version 4.3.2. This method was also applied to examine the correlation between PICs of weighted average gene family sizes and genome sizes.

### Classification of Faster-evolving and Slower-evolving Lineages

To examine the variation in the weighted average gene family size within 12 orders, we utilized the R package diptest version 0.76.0 for conducting unimodality tests separately on the evolutionary rates (measured as the branch length from the tip to the Saccharomycotina common ancestor (SCA) on the phylogenetic tree) and the weighted average gene family sizes for each of the 12 orders. Additionally, we applied the same method to analyze the branch length from the tip to the most recent common ancestor of the order in focus, which yielded the same results. For orders exhibiting significant non-unimodal distributions in both evolutionary rates and weighted average gene family sizes, we applied density-based spatial clustering of applications with noise (DBSCAN) algorithm using the R package dbscan version 1.1.12 to identify clusters based on evolutionary rates. Additionally, we mapped weighted average gene family sizes onto the phylogenetic tree to examine lineage-specific variations. In orders displaying lineage-specific variations, the DBSCAN clusters with faster evolutionary rates were labeled as faster-evolving lineages (FELs), and those with slower rates were identified as slower-evolving lineages (SELs).

### Analysis of Gene Family Expansion and Contraction Between Faster and Slower Evolving Lineages

To determine which gene families exhibited expansion or contraction in FELs compared to their SEL relatives, we performed a fold change analysis using a custom R script, based on the method developed in the previous study^1^. For a given yeast order with FEL and SEL lineages, we first calculated the average copy numbers (non-weighted) for each gene family within the FELs and SELs, respectively, then divided the average value of FELs by that of SELs. Additionally, we performed the Kolmogorov-Smirnov (KS) test using the ks.test function from the R package stats version 4.3.2, coupled with the Bonferroni method for p-value adjustment, to ascertain the significance of these expansions or contractions. Consistent with the criteria established in prior research^1^, we reported those gene families that underwent significant changes (adjusted *P* ≤ 0.05), and a fold change exceeding 1.5 for expansions or less than 0.67 for contractions. A fold change of 0 was interpreted as a loss of the gene family, while a fold change nearing positive infinity indicated the acquisition of a gene family.

### Principal Component Analysis of Gene Family Presence and Absence Pattern

To compare the difference of gene family composition across yeasts, we conducted a Principal Component Analysis (PCA) based on the presence (1) or absence (0) data of gene families^3^. We first discerned conserved and species-specific gene families by setting average coverage threshold at 0.1 based on the density plot (Figure S1). Gene families with the average coverage equal to or exceeding 0.1 were considered conserved, while those below the threshold were classified as species-specific. We employed PCA on both conserved and species-specific gene families using the R package stats version 4.3.2. Consequently, we performed density clustering to the PCA results using the dbscan function from the R package dbscan version 1.1.12, grouping species with similar distribution patterns into distinct clusters. We also conducted the same analysis using a more stringent threshold of 0.5 in the PCA to exclude more noise from species-specific and/or rare gene families, which yielded consistent results.

To identify key gene families driving the distribution of yeasts along the first or second principal components, we employed a custom R script for detailed analysis (Figure S12). Initially, we ranked gene families according to their contribution (from the rotation table in the PCA results using the R package stats) to each principal component (PC), both in ascending and descending order. To identify the optimal number of top-ranking gene families whose average presence values best correlate the coordinates of yeasts, we calculated the average presence values for the top 1, 2, i, and up to top n gene families (where i is the specific number of gene families, and n is the total number of gene families). The average presence value for the top n gene families was determined by dividing the total presence of these n gene families in a species by n. Subsequently, we conducted a Spearman correlation test using the cor.test function (method = spearman) from the R package stats version 4.3.2. This test assessed the relationship between the average presence values of species in the top i gene families and their respective positions on the PC. The gene families with the highest absolute correlation values were selected. A positive correlation indicates that species with larger coordinates on the PC tend to have more gene copies in the top i gene families, while a negative correlation suggests that species with larger coordinates are likely to have fewer copies of these gene families.

### Analysis of the Rates of Speciation and Carbon Source Utilization Trait Gain and Loss

To investigate whether different carbon source utilization traits are more readily acquired or lost in the FELs or SELs, we used the analytical method and carbon source utilization data from previous studies^11^. Firstly, we pruned the species tree to only retain yeasts with available metabolic data, resulting in trees comprising exclusively Dipodascales FEL or SEL species. Subsequently, we employed BayesTraits version 4.0.0 and its reverse jump model^63^ to conduct two simulations for each carbon source. The first simulation set the loss rate of carbon source utilization traits equal to the acquisition rate (using the parameter “Res q01 q10”), while the second did not equate these rates (no specific parameter used). Additionally, each model underwent 10,100,000 iterations, using 200 stepping stones, with sampling every 1,000 iterations. The burn-in was set at 100,000 iterations. We also employed the R package coda version 0.19.4 for visualization purposes to ensure model convergence.

To select the appropriate model for determining whether the loss rate of carbon source utilization traits should be equal to or different from the acquisition rate, we calculated Log Bayes Factors according to the BayesTraits manual (https://www.evolution.reading.ac.uk/BayesTraitsV4.1.1/BayesTraitsV4.1.1.html). Log Bayes Factors is utilized to compare the relative evidence between two statistical models. When the Log Bayes Factor is less than 2, we opt for the relatively simpler model (where the acquisition rate is equal to the loss rate). Conversely, when it is 2 or higher, we select the more complex model (where the acquisition rate is not equal to the loss rate). Subsequently, based on the selected model, we count the number of instances where the carbon source utilization trait’s acquisition rate is either greater than or less than its loss rate. If the instances of the acquisition rate being higher than the loss rate significantly outnumber those where it is lower, we conclude that the lineage tends to acquire that particular carbon source utilization trait. On the other hand, if there are more instances of the acquisition rate being lower than the loss rate, the lineage is considered more inclined to lose that trait. If the simpler model is chosen based on the Log Bayes Factor, we infer that the lineage is neither inclined to lose nor to acquire the carbon source utilization trait.

To investigate the connections among gene family expansions and contractions, the acquisition and loss of carbon traits, and the diversification of species, we estimated speciation rates from the DR statistic^64,65^ calculated using the inverse equal splits method^66^ using a recently published time-calibrated phylogeny^11^.

### Investigation of Metabolic Pathways, the Spliceosome Pathway, and the DASH Complex

To investigate how gene loss might affect crucial biological processes in the FEL of Dipodascales, we used *S. cerevisiae* as a reference to map gene names to its pre-mRNA splicing pathway, metabolic pathways, and the DASH complex. We first identified gene families that exhibited significant contraction or loss in the fold change analysis and those that were representative in contributing to the principal component, using the representative genes from *S. cerevisiae*. If an *S. cerevisiae* gene was assigned to a gene family according to OrthoFinder results, we named the gene family using the *S. cerevisiae* gene name. Otherwise, the gene family remained unnamed due to the uncertainty of its classification. Subsequently, we used these gene family names for pathway mapping. Specifically, we used the search function on the KEGG website (https://www.genome.jp/pathway/sce03040) for the pre-mRNA splicing pathway, the Highlight Gene(s) feature on the *Saccharomyces* Genome Database (SGD)^28^ biochemical pathways site (https://pathway.yeastgenome.org/overviewsWeb/celOv.shtml) for metabolic pathways, and the previous study^31^ for DASH complex.

To verify the absence of genes indicated in our gene copy numbers heatmap (Figure 4a), we carried out independent orthology delineation using InParanoid version 4.2^67^ and sequence search using Basic Local Alignment Search Tool (BLAST) version 2.15.0+^68^. This was to ensure accuracy and address potential misassignments by OrthoFinder or errors in genome annotations. With InParanoid, we compared the protein-coding genes from all species in our heatmap against those of *S. cerevisiae* to identify orthologous genes, confirming that species depicted as lacking certain genes genuinely did not have those orthologs. We also used blastp (e-value threshold of 1e-5) to compare species, which are shown as missing gene families in the heatmap, with a reference species that contained all genes in our heatmap. For addressing potential annotation inaccuracies, we performed genome-protein comparisons using tblastn (also with an e-value of 1e-5).

### CAFE Analysis of Gene Copy Number Evolution

To estimate gene family expansion and contraction events, we utilized computational analysis of gene family evolution using (CAFE) version 5.0^69^. Due to the computational limitation of CAFE in processing the complete analysis of 1,154 genomes, we first employed separate analyses for each of the 12 orders. For these analyses, the input time tree was pruned to include only species from the order under study. The input gene families needed to meet any of the three criteria: 1) presence in the MRCA of studied order, as determined by maximum parsimony; 2) presence in the studied order and at least one of the remaining 11 orders; and 3) presence in both the studied order and the outgroup. Gene families not meeting those three criteria are specific to the order under study, and thus are irrelevant for the CAFE analysis of 12 order MRCAs. The estimated gene contents of each yeast order were then analyzed by CAFE to reconstruct gene family copy numbers at the SCA. The input time tree was pruned to only include the MRCAs of each of the 12 yeast orders.

We experimented with different numbers of gamma categories (k ∈ [2, 10]) using the “-k” parameter and selected the k value with the highest likelihood. To determine the alpha (the evolutionary rate of genes within gene families over time) and lambda (the rate of increase or decrease of gene families over time) values, we ran 10 iterations with the determined k value and chose the alpha and lambda values that yielded the maximum likelihood.

To confirm the reliability of our CAFE analysis on the full dataset of 1,154 species, we used the same methods on subsampled datasets of 200 species and 50 species. The 200 and 50 species datasets were subsampled based on genome completeness from BUSCO results, ensuring that at least one species from each order was included.

To ensure robust reconstruction of ancestral node gene contents, we only displayed gene families that met any of the following criteria: 1) present in the SCA, as determined by maximum parsimony; and 2) present in only a specific order and the MRCA of that order.

### Orphan Gene Families

We defined orphan gene families as those specific to a particular order and exhibiting a high species coverage within that order. Specifically, an orphan gene family is characterized by being present in at least 98%^26^ of the species within a given order. This means that to qualify as an orphan, a gene family must be found in 98% or more of the species within the order under consideration. Additionally, these gene families must be completely absent in all other remaining orders.

### Functional Enrichment Analysis

We conducted functional enrichment analyses of gene families across fold change, PCA, and CAFE analyses. For the fold change analysis, the background set for enrichment consisted of the union of gene families present in all yeast species within the studied order. In PCA, particularly for the top 610 gene families linked with PC1, the background was composed of all gene families involved in the PCA. For the CAFE analysis, the background for enrichment was the set of gene families included in the gene family copy number table used as input. Our enrichment analyses drew upon various annotations, including GO annotations, KEGG annotations, and InterPro annotations. The correspondence description tables for GO terms and KOs and InterPro entries were downloaded from the GO (https://geneontology.org/), KEGG (https://www.genome.jp/kegg/) and InterPro (https://www.ebi.ac.uk/interpro/) websites, respectively, on November 23, 2023.

All enrichment analyses were conducted using the R package clusterProfiler version 4.6.0^70^ with default parameters, selecting only results with *P* ≤ 0.05. To translate GO terms into more generalized and concise GO slims in fold change enrichment analysis, we employed GOATOOLS version 1.2.3^71^. For this process, we utilized the go-basic.obo and goslim_yeast.obo files, which were retrieved from the Gene Ontology website on December 13, 2023.

### Data Visualization

We utilized the R package ggtree version 3.8.0^72^ to visualize phylogenetic trees and associated CAFE data, and ggplot2 version 3.4.3 for other graphs. Images representing taxa were hand-drawn, sourced from PhyloPic (https://www.phylopic.org/), and customized in terms of color using rphylopic version 1.3.0.

### Data and Code Availability

The reference phylogeny of yeasts, along with genome and annotation data for yeasts, Pezizomycotina, animals, and plants, are accessible from previous studies described above. Additionally, NCBI taxonomy and source details for this study can be found in Table S1. We have deposited all new functional annotations, analyses, and codes in the Figshare repository.

## Acknowledgments

We thank Li Lab and Y1000+ members for helpful discussions. This work is supported by the Ocean Negative Carbon Emissions (ONCE) Program. Y.L. was supported by the National Key R&D Program of China (2023YFA0915501), National Natural Science Foundation of China (No. 42376147), and Shandong University Outstanding Youth Fund (62420082260514). X.Z. was supported by grants from the Basic and Applied Basic Research Foundation of Guangdong Province (2022A1515010223) and the National Natural Science Foundation of China (No. 32260652). J.L.S. is a Howard Hughes Medical Institute Awardee of the Life Sciences Research Foundation. Research in the Hittinger Lab is funded by the National Science Foundation (DEB-2110403), USDA National Institute of Food and Agriculture (Hatch Project 7005101), in part by the DOE Great Lakes Bioenergy Research Center (DOE BER Office of Science DE–SC0018409), and an H. I. Romnes Faculty Fellowship (Office of the Vice Chancellor for Research and Graduate Education with funding from the Wisconsin Alumni Research Foundation). Research in the Rokas lab is supported by the National Science Foundation (DEB-2110404), NIH/National Institute of Allergy and Infectious Diseases (R01 AI153356), and the Burroughs Wellcome Fund.

